# Interacting roles of lateral and medial Orbitofrontal cortex in decision-making and learning : A system-level computational model

**DOI:** 10.1101/867515

**Authors:** Bhargav Teja Nallapu, Frédéric Alexandre

**Author notes:** 200 Avenue de la Vieille Tour, 33405 Talence, France. (FA). BTN is the graduate student who implemented the model, ran the simulations, and wrote the article. FA corrected the article. BTN and FA together conceived the idea and equally contributed to the conception and discussion.

## Abstract

In the context of flexible and adaptive animal behavior, the orbitofrontal cortex (OFC) is found to be one of the crucial regions in the prefrontal cortex (PFC) influencing the downstream processes of decision-making and learning in the sub-cortical regions. Although OFC has been implicated to be important in a variety of related behavioral processes, the exact mechanisms are unclear, through which the OFC encodes or processes information related to decision-making and learning. Here, we propose a systems-level view of the OFC, positioning it at the nexus of sub-cortical systems and other prefrontal regions. Particularly we focus on one of the most recent implications of neuroscientific evidences regarding the OFC - possible functional dissociation between two of its sub-regions : lateral and medial. We present a system-level computational model of decision-making and learning involving the two sub-regions taking into account their individual roles as commonly implicated in neuroscientific studies. We emphasize on the role of the interactions between the sub-regions within the OFC as well as the role of other sub-cortical structures which form a network with them. We leverage well-known computational architecture of thalamo-cortical basal ganglia loops, accounting for recent experimental findings on monkeys with lateral and medial OFC lesions, performing a 3-arm bandit task. First we replicate the seemingly dissociate effects of lesions to lateral and medial OFC during decision-making as a function of value-difference of the presented options. Further we demonstrate and argue that such an effect is not necessarily due to the dissociate roles of both the subregions, but rather a result of complex temporal dynamics between the interacting networks in which they are involved.

**Author summary:** We first highlight the role of the Orbitofrontal Cortex (OFC) in value-based decision making and goal-directed behavior in primates. We establish the position of OFC at the intersection of cortical mechanisms and *thalamo-basal ganglial* circuits. In order to understand possible mechanisms through which the OFC exerts emotional control over behavior, among several other possibilities, we consider the case of dissociate roles of two of its topographical subregions - *lateral* and *medial* parts of OFC. We gather predominant roles of each of these sub-regions as suggested by numerous experimental evidences in the form of a system-level computational model that is based on existing neuronal architectures. We argue that besides possible dissociation, there could be possible interaction of these sub-regions within themselves and through other sub-cortical structures, in distinct mechanisms of choice and learning. The computational framework described accounts for experimental data and can be extended to more comprehensive detail of representations required to understand the processes of decision-making, learning and the role of OFC and subsequently the regions of prefrontal cortex in general.

## Introduction

Psychological and economic accounts of human and animal decision making have placed a great emphasis in the concept of value [1–3]. Value-based decision making is a process of evaluation of alternatives in terms of subjective preference to their potential consequences and their relevance to internal motivations. Thus often in the context of decision-making of an animal, *value* remains a subjective, comparable quantity, that spans across multiple neural pathways [4]. The Orbitofrontal Cortex (OFC) is one of the prominent regions of the Prefrontal Cortex (PFC) that is believed to play a crucial role in value-based decision making [5, 6] and learning [7] where *value* changes over time because of changes in internal states or external contingencies. However, specific functional role of the OFC in value-based decision making is not clear. Moreover, to some degree or the other, the OFC has been implicated in almost all of the fundamental processes formally described to be involved in value-based decision making [8] - Representation, Valuation, Action selection, Outcome evaluation and Learning.

In the context of ‘Representation’ of a decision scenario, the OFC has been proposed to represent a cognitive map of task space [9]. Further in the context of ‘Valuation’, the role of OFC has been implied in encoding the *value* of the offered and chosen goods [10, 11]. It might appear that there are other valuation systems in the brain [12–14], but what makes OFC unique in valuation is that it encodes the value irrespective of the visuo-spatial and motor aspects. For instance, the value representation in the OFC has been argued, not only to guide choices consistent with *Transitivity* [15], but also to represent largely varying subjective values in an adaptive manner [16, 17]. These representations were proposed to be based on a *common currency* [1, 6, 18] that guide the comparison for the decision between different objects that are otherwise incomparable. Alternative to the theory of *common currency*, it was proposed that what the OFC facilitates is the process of *common scaling* [19–21] which is qualitatively distinct from that of converting different rewards into a *common currency*. Instead, *common scaling* corresponds to retaining the individual value of each reward, and converting them to a different scale that makes them comparable. Evidently, with complex possibilities in the process of valuation, arise different possibilities of action selection processes.

Furthermore, exact representations and mechanisms through which the OFC contributes to behavior are still up to active debate [22]. Moreover, several early implications of the OFC in the paradigms related to reversal learning [23], response inhibition [24, 25], flexible stimulus-outcome associations [26, 27] have been overturned using the same experimental techniques [7] or even more accurate ones [28], modified task structures [29, 30] or pointing out the fact that the findings from other related brain regions explain certain implications better [31–35].

### Dissociate roles of lateral and medial OFC

The evident underlying complexity of studying the role of the OFC in value-based decision making and learning, and goal-directed behavior is underlined by the large heterogeneity of the region, unlike the rest of PFC which is homogeneously granular. The heterogeneity is multi-fold : different groups of neurons that encode different aspects of choice process in a single task context [11], cyto-architecturally different areas (granular and agranular) and their remarkably distinct connectivity pathways through different brain structures [36–38]. The possibility of functionally dissociate roles of topologically different sub-regions of the OFC has been of wide interest recently. While there are other sub-divisions of the OFC reported to be playing functionally distinct roles in behavior [39–41], the distinction that is most extensively reported to imply strikingly different functional roles is the one between lateral and medial parts of OFC [28, 42, 43]. In the scope of this article, as referred in most of the related experimental works, ventromedial prefrontal cortex (vmPFC) is also considered under the purview of medial OFC [44, 45]. It has also been observed that lateral and medial OFC have clear divergent connections to different networks [46]. Both in monkeys and humans lateral OFC is reported to receive extensive projections from diverse sensory modalities through the somatosensory and insular cortices, and also heavy projections from amygdala [47–49]. Whereas, the medial OFC has strong projections from hippocampus, hypothalamus, ventral striatum (VS), relatively less projections from amygdala, and is strongly connected with the cingulate cortical areas [50, 51].

In this current work, we present a recurrent neural network model of decision making and learning involving the OFC. The OFC, together with some nuclei of the basal ganglia (BG)(especially VS) and the thalamus (Th), forms a closed loop whose dynamics leads to action selection by competition resolution. This loop is a part of several similar generic loops that are formed between different cortical regions and different nuclei of the BG. A generic loop will be referred hereafter as a CBG loop. Notably, we separate the part involving the OFC into two CBG loops involving lateral and medial OFC, accounting for the individual experimental implications of the lateral and medial sub-regions. The input to the lateral OFC loop is provided with the information that represents *exteroception* - the value information arising from external factors like visual cues; the input to the medial OFC loop is predominantly the *interoception* - value information more with respect to internal motivational processes like satiety levels and internal needs [42]^1^. Across both these loops, we use the idea of the Current Subjective Value (CSV) in the model as an input to lOFC and mOFC. Besides the activation in the loops that represents the visual salience of the cues, CSV represents the value based on which the sub-regions of OFC contribute to the decision-making process. Such a subjective value is known to arise from a comprehensive relation of lateral OFC with basolateral amygdala and the ventral striatum in an ongoing task context [52–55] (see Materials, CSV).

We provide a plausible explanation for one of the prominent experimental observations regarding the dissociate roles of lateral and medial OFC, studied in individual lesions in monkeys [43]. We represent this proposed dissociation in terms of the representation and processing of the task information. Furthermore we argue that, in the context of *learning* vs *choice* notion of lateral and medial OFC [56], more than a clear dissociation, it is the temporal interaction of both the sub-regions that highlights their roles at different stages of a decision task.

## Results

We first describe the performance of an existing model of decision-making and learning on a 2-arm bandit task with probabilistic reward. We take the advantage of generic nature of the task to highlight the fundamental dynamics of the model. We then show that the model presented here with the distinct description of lateral and medial OFC replicates the results of basic model, robustly and in more realistic timescales. We further present complementary findings of separate lesions (simulated) of the lateral and medial OFC components in the model. We discuss the effect of these findings on the performance in different task contingencies, replicating a neuroscientific evidence found in monkeys with lesions to different subregions of OFC.

### 2-Arm Bandit Task and Probabilistic Reward Learning

Multi-arm bandit task is a classic reinforcement learning problem that has been used in the study of decision-making in experimental [7, 43, 57] and computational neuroscience [58–60]. Typically, in an N-arm bandit task, there are *N* possible cues (bandits) each carrying a different probability of reward and requiring a particular action to do, in order to select the cue. Fig. 1A shows an example trial of a 2-arm bandit task that has been used to study the computational models of probabilistic reward-based learning involving the basal ganglia (BG) [60, 61]. In this case, cue is one of the four possible shapes. The reinforcement in the model during the task is driven by the probabilistic reward offered at the end of each trial, with a different probability for each cue. It has been shown that monkeys learn to perform the task [57], learning the reward contingencies over time and choosing always the best rewarding option after learning.

**Fig 1.**
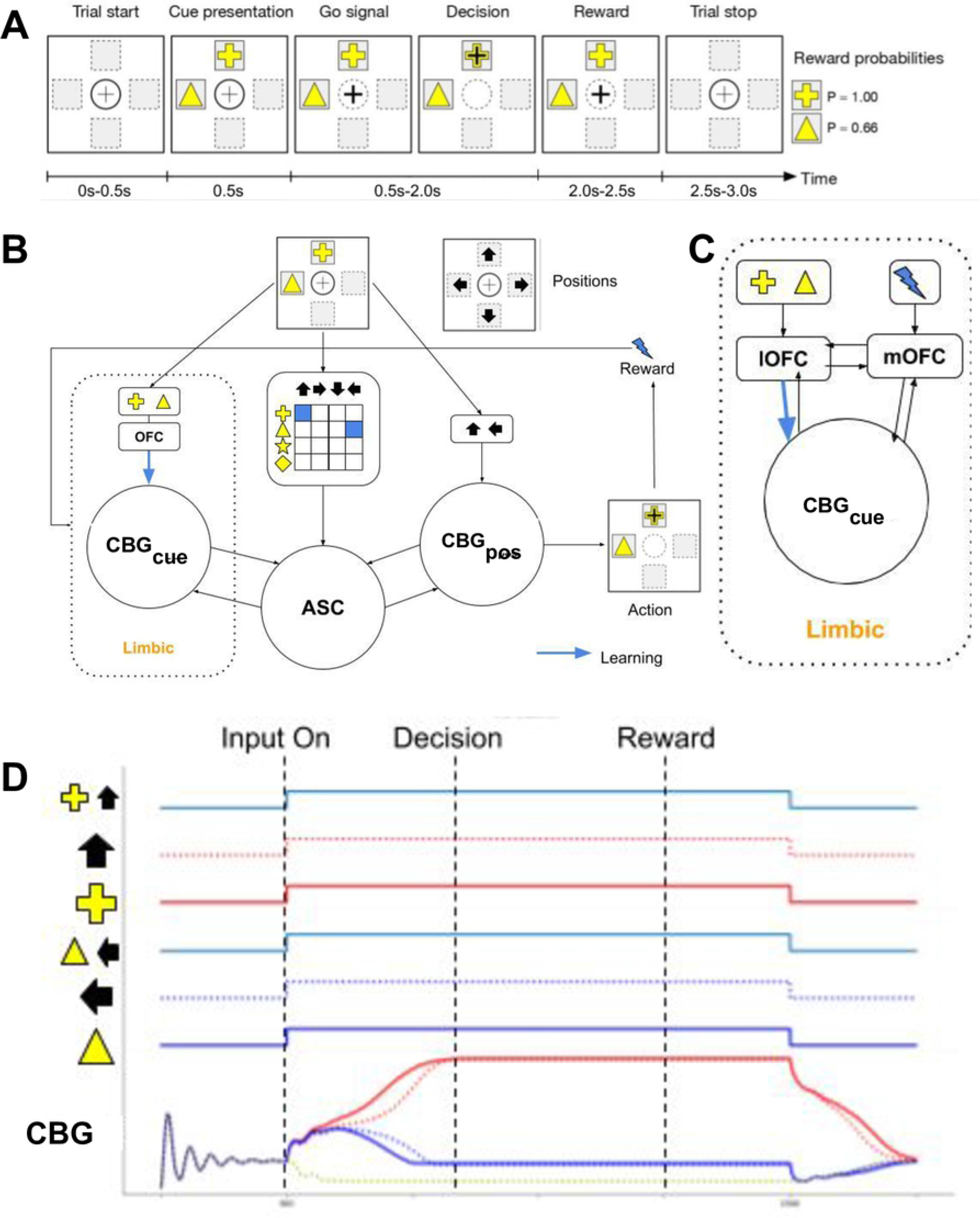
2-arm bandit task. **A.** Sample trial from a 2-arm bandit task. Out of four possible shapes (cues) are shown at two random positions (of the four cardinal positions). The position that is chosen implies the choice of the shape made. **B.** Basic model involving two CBG loops and an associative loop (ASC), one CBG loop leading to a choice between the two cues and the other between the two positions. The final output that is considered from the model within a trial is that of the decision of CBG position, the cue shown at the *chosen position* is considered as the *chosen cue*. Note the CBG Cue is labelled *limbic*, as it will be developed more into components representing sub-regions of the OFC. Blue arrow represents the connection that can be modified by learning. **C.** The proposed change in the original model which will be described in detail in the following section. **D.** Activation of each cue that is shown in a choice, its position and the combined information. Also, the evolution of activity in a CBG loop - solid lines for cue, dashed lines for positions.

The basic model (referred hereafter as OFC model) is a set of inter-connected CBG loops and an associative network (ASC), each network processing different information and contributing for a decision within the network (Fig 1C). In each trial, the CBG_*cue*_ labeled ‘limbic’ takes as the input, the activation for the shapes that are presented in the trial. This activation represents a constant visual salience component, that in the simplest case, is same for every stimulus (shape). Similarly the other CBG position loops (CBG_*pos*_) takes as the input, the activation of the positions where the shapes are presented. Since the positions are chosen randomly and carry no significance in obtaining reward, there is no value-learning in this CBG_*pos*_ loop. Hence the activation of a position represents just the presence of a cue at that position. Finally, the ASC network takes as the input, the combined information of binding specific shape to a specific position. The ASC network represent the associative loop through lateral PFC and the dorsomedial striatum (DMS) which is believed to represent a multi-modal information of stimulus-vs-position mapping [62]. This is implemented in the form a 2 dimensional mapping for each shape against all possible position and each position against all possible shapes (Fig 1B, blue squares). The networks are inter-connected in such a way that while each of the CBG loops independently processes the information that it is activated with, it also affects the activities in the other through the ASC network. The network architecture within each CBG loop that guarantees the resolution of competition between the options is based on classical BG pathways that have been previously explained with computational accounts [59, 60, 63].

In each trial of the task, the model is presented with pseudo-randomized pairwise presentation of the four possible shapes in any two of the four possible positions (’Cue presentation’ phase in Fig 1A and first 6 panels in Fig 1D). Although the performance of the model is assessed in terms of the shape it chooses for optimal reward probability, the choice is confirmed only if the corresponding position of the shape is chosen as the ‘motor’ decision (Fig 1A, black + sign under ‘Decision’ phase, dashed lines under ‘CBG’ in Fig 1D). Thus, after the ‘Decision’ phase of the trial, the shape at the chosen position is considered as the choice of ‘cue’ and the reward is delivered according to the predetermined probability associated to that cue.

The performance of the model is demonstrated under two conditions : EASY and DIFFICULT. EASY is the condition where the reward probabilities related to each shape are fairly separated and DIFFICULT is the condition where the reward probabilities are either lower or closer, thus making the reinforcement difficult (Fig 2A). The effect of learning in the model after each trial can be observed in terms of the decision times over the duration of the task. A decrease in decision times of both cue and position is observed (Fig 2B, left). A running average over the choice of 10 trials is considered for the performance over 120 trials. The performance of the model under the EASY condition replicates animals’ behavior [57] (Fig 2D, blue). In the DIFFICULT condition (Fig. 2A, right), the reward probabilities of both the shapes are lower or closer. This should result in lower rate of reinforcement and thereby make it difficult to make a correct choice. Animals however, with considerable amount of training, were shown to identify the option with more chance of reward and thus make correct choices [7, 43]. We tested the same model as in the previous EASY case (Fig 2A, left), but the model couldn’t learn the appropriate contingencies well. The Decision Times (DTs) were longer compared to the previous case (Fig 2C) and the overall performance was sub-optimal (Fig 2D, red).

**Fig 2.**
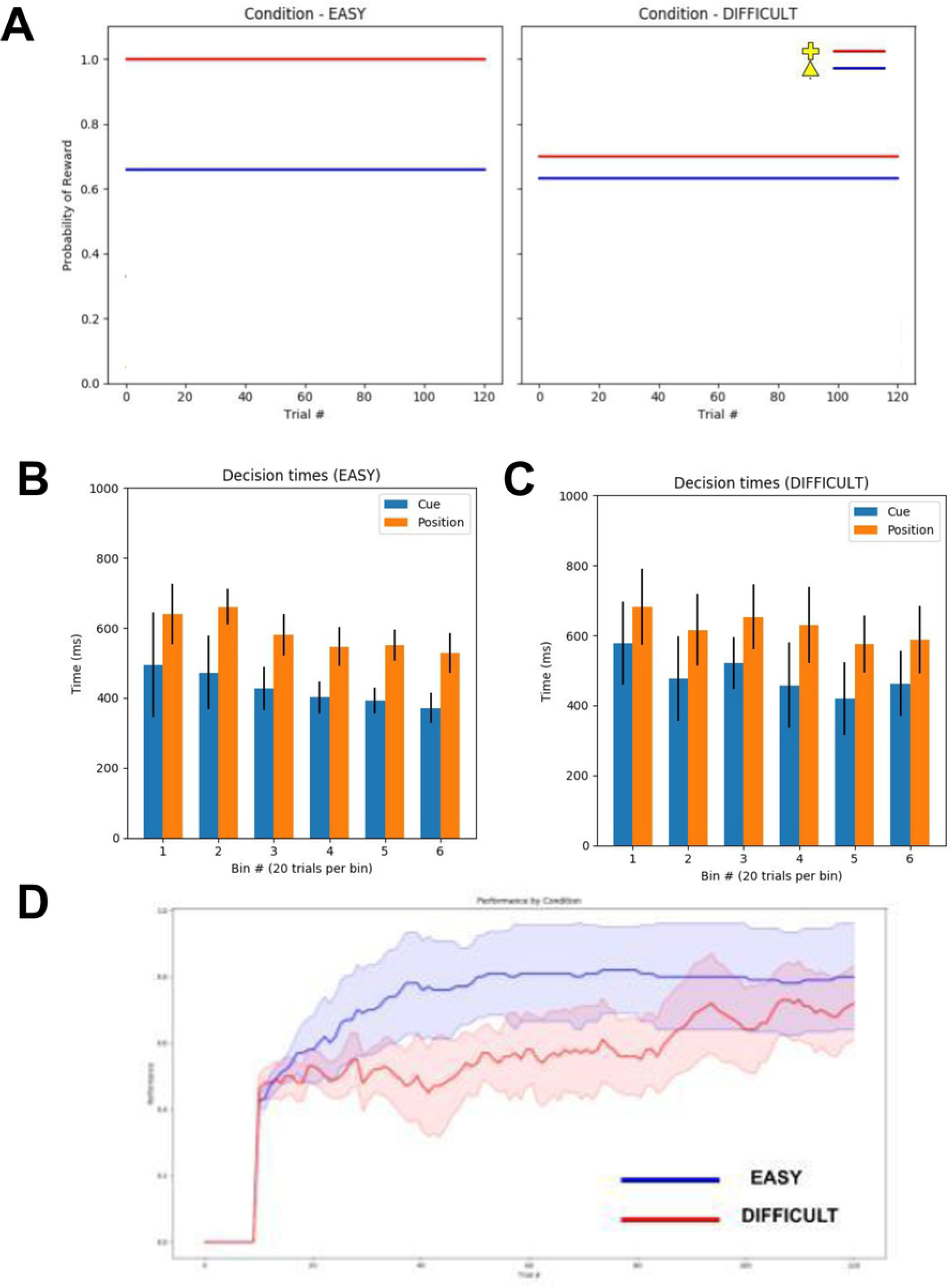
N-arm bandit task. The task described in Fig 1**A** has two possible cues (shapes) each with a predetermined probability of reward upon choice. **A.** Left and right figures show two different reward probability schemes, in EASY and DIFFICULT task scenarios. Each color represents a particular shape, as in the legend of right sub-figure. **B, C.** Decision Times (DTs) in the model after cue presentation. 120 trials are divided into 6 bins, with 20 trials per bin, and the DTs of both cue decisions and the position decisions are averaged per bin. **B** shows the DTs in EASY condition of the task and **C** in DIFFICULT condition. **D.** Performance of the model. Running average of number of correct choices across 10 trials, averaged over 10 sessions. Correct choice means the shape that rewards the most according to the predetermined probabilities. Lighter color filling represents the standard deviation.

### Precise Value Comparison

We then extend the ‘limbic’ CBG loop to individually describe two separate CBG loops - one representing the lateral OFC and the other representing the medial OFC. Here after this version of the model will be referred as *lmOFC* model. The CBG loop involving lateral OFC builds on the top of the single limbic loop from the basic model (described in Fig 1B). In addition to the activation (*I_ext_*) to the network, a Current Subjective Value (CSV) for each shape is also added to the input. CSV represents the subjective value of a shape at any moment taking the externally learned reward contingencies and internal bodily desire for the reward that the shape leads to (see *Materials*, CSV). Another key aspect of lOFC is that it properly assigns the obtained reward to the appropriate choice made in that trial (referred as *credit assignment*). There has been evidence that neurons in lateral OFC are particularly active after the reward delivery in a choice [42] and also the fact that medium spiny neurons neurons which are extensively involved in decision-making are consistently active for a while after reward delivery [55]. These evidences support the possibility that cortico-striatal synaptic plasticity is a plausible phenomenon in the context of obtaining reward. Similar arguments were made by other experimental findings [7].

The CBG loop with medial OFC receives input from the CSV layer. Medial OFC has a separate value comparison mechanism implemented as a simple ‘recurrent excitation lateral inhibition’ model, activated by the CSVs received. It was shown that the activity in medial OFC correlated to the value difference between the options [64]. Supporting the view that the relative difference of the presented options is represented in vmPFC, multiple value comparison mechanisms have been proposed. This value difference signal further allows vmPFC to perform a value comparison to facilitate the choice through principles of recurrent excitation and lateral inhibition [21, 65–67]. The output activities of mOFC are fed into its CBG loop. It has been shown that one of the general function of populations in the PFC is to maintain history of decision events such as previous action, previous reward etc [68]. Accordingly, we implemented a simple history of rewards in mOFC, without cue-specific information. As the lOFC maintains the current choice until the reward delivery and later [42], possibly a history of choices is maintained in lOFC. It was shown that lesions to lOFC affect the appropriate consolidation of the reward history with the choice history [7]. Hence, for the sake of simplicity, both the histories in lOFC and mOFC are combined within the lOFC to provide a combined choice-reward history up to one previous choice and reward, and it is fed into the CBG loop of lOFC along with the activation of the cue. In addition, a synaptic connection is added to the ASC layer outside the limbic network, from each cue population in lOFC to all the possible position populations in the 2-D mapping of ASC network. However the learning in these connections would be less influential on the decision, compared to that in the lOFC network, since the learning happens with respect to all four possible positions corresponding to the cue in the 2-D mapping of ASC (Fig 1B, input to ASC). Although it has been shown that mOFC / vmPFC encodes action-outcome associations in several task settings [69–72], the design of the n-arm bandit task setting does not allow much of learning action-values. The reason is that the task randomizes the positions where the cues are present and hence the action required to chose a cue.

We then tested the lmOFC model on the DIFFICULT condition as in the previous task. The model performed considerably well compared to the previous OFC model, with much faster DTs. Both the models have an estimated value difference for the ongoing task, across all the trials. Interestingly, the precise value comparison in mOFC estimates the value difference across all the trials better than that estimated by the OFC model under DIFFICULT condition (Fig 3D).

**Fig 3.**
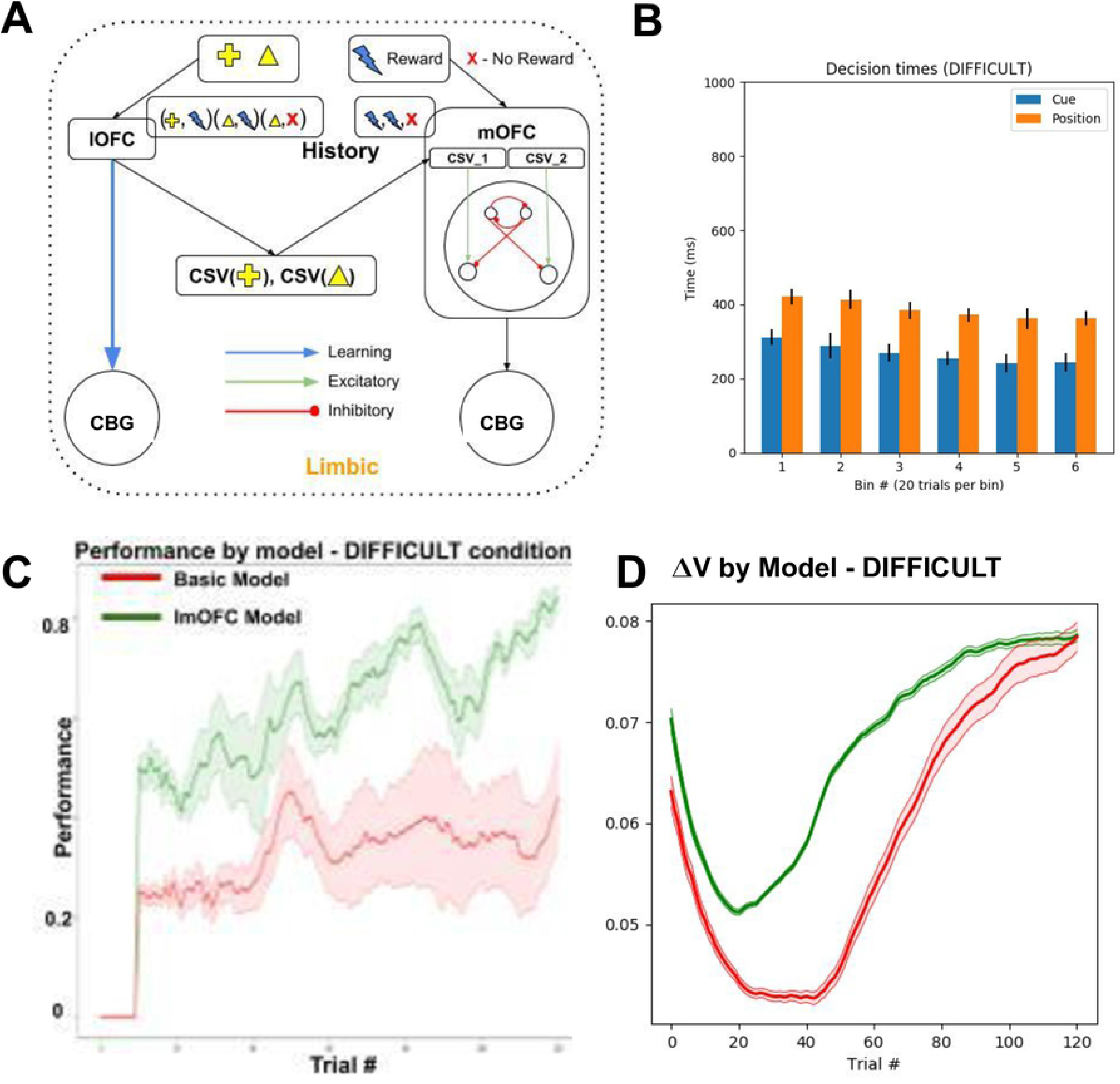
lmOFC Model : CBG loops with lateral and medial OFC. **A.** lmOFC Model. Changes in the ‘limbic’ CBG loop, compared to the basic model. Lateral OFC (lOFC) has access to cue identity (shape), hence drives learning the connections to its CBG loop. lOFC also activates the Current Subjective Value (CSV) for each of the presented cues from elsewhere. Medial OFC (mOFC) has a value comparison mechanism to compare the CSVs of the presented cues it receives. mOFC further drives its CBG loop with the ongoing value comparison outputs. Both lOFC and mOFC also maintain general history of chosen cue-reward association and reward respectively. This input is also used in the activation to their respective loops. **B.** The average DTs of decisions choosing cue and position, across 120 trials binned every 20 trials. **C.** The performance of the lmOFC model (green) in comparison with the performance of the basic model in DIFFICULT condition (red). **D.** Average value difference of the presented options estimated in lmOFC model (green) and basic model under DIFFICULT condition (red).

### Proximity of Values and Decision Making

We tested the lmOFC model on a 3-arm bandit task (Fig 4). Each of the three cues that are shown in every trial has a reward probability upon its choice. As shown in Fig 4, V1, V2 and V3 are the reward probabilities associated to the cues *plus*, *delta* and *star* respectively in a given experimental session. The task is carried out under three different reward schedules (Fig 5A-C). In all the sessions, V1 and V3 are fixed to be .7 and 0.05. V2 value is changed across three types of sessions : V2_HIGH, V2_MID and V2_LOW where V2 is set to 0.6, 0.3 and 0.1. Similar task schedule was used on animals to test the effects of lesions of lateral and medial OFC separately [43].

**Fig 4.**
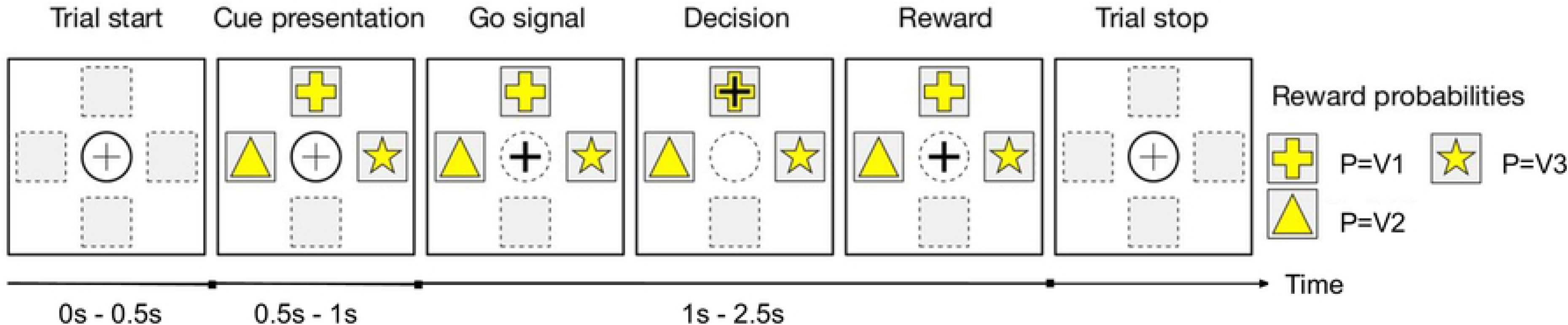
3-arm bandit task. A sample trial from the 3-arm bandit task. Three possible shapes (cues) are shown in three random positions (of the four cardinal positions). The position that is chosen implies the choice of the shape made. Upon selection of a shape, a reward is delivered with a probability (p), which is different for each of the shapes (V1, V2 and V3).

**Fig 5.**
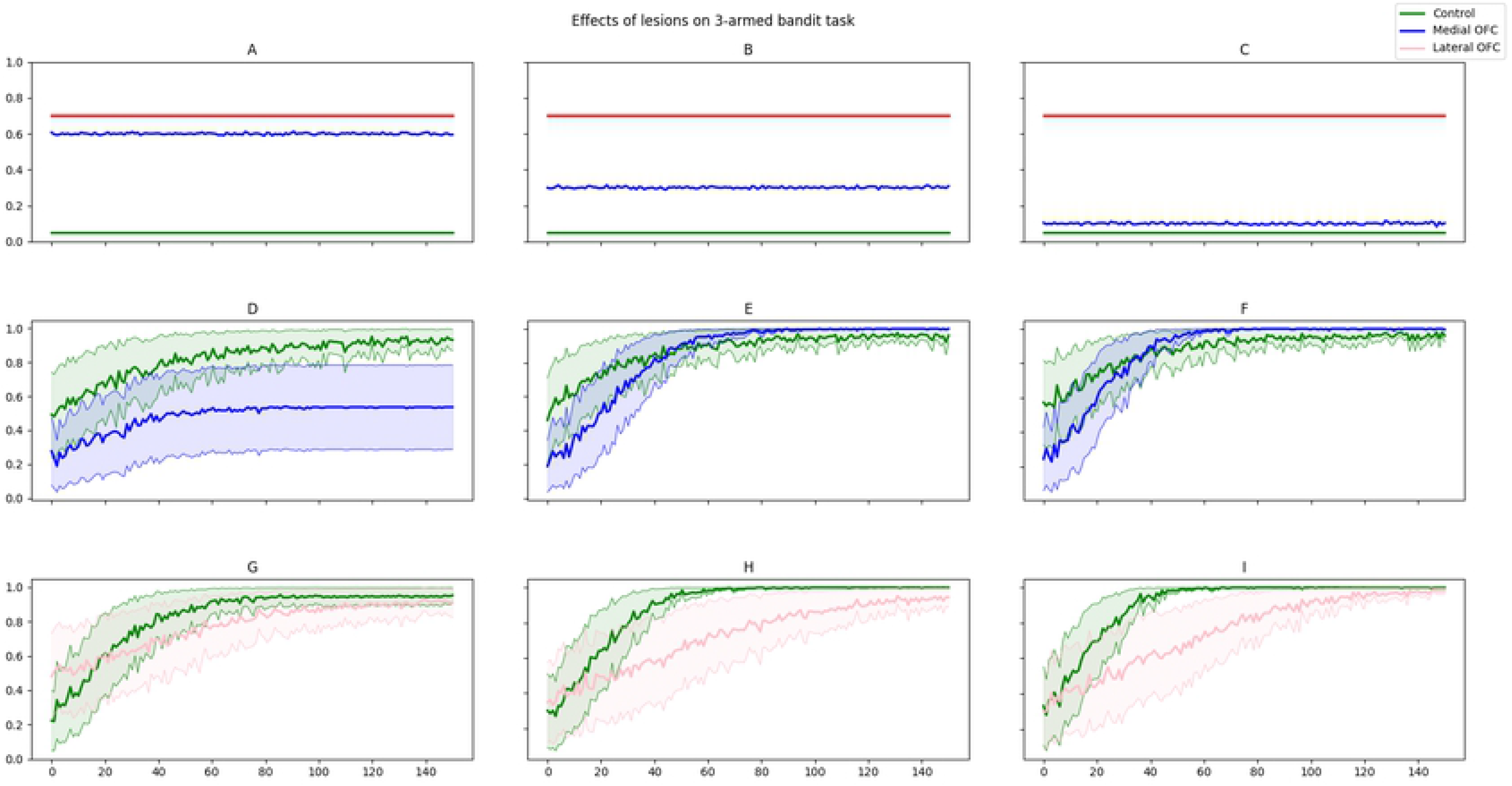
Effects of lateral and medial OFC lesions in the model. **A-C.** 3 conditions of the task (each column). Of the reward probabilities V1, V2 and V3 described in Fig 4, V1 and V3 are fixed in all 3 task conditions (A-C, red and green respectively). The 3 conditions depend on the value V2 (A-C, blue) : V2_HIGH, V2_MID and V2_LOW. **D-F.** Lesion of mOFC under each task condition. The average performance in each condition with a lesion to mOFC (blue) is compared to the control performance (green). **G-I.** Lesion of lOFC under each task condition. The average performance in each condition with a lesion to lOFC (pink) is compared to the control performance (green).

The lmOFC model is used without any changes. At the time of presentation in every trial, 3 cues are activated simultaneously (along with their positions in the CBG_*pos*_ loop and in the ASC network). In terms of the model parameters, just the input activation which represents the cue salience had to be increased as compared to when the choice was between 2 options (as in the previous tasks). In this task, a correct choice or a good choice is a V1 choice. The model reached optimal performance (more than 80% V1 choices) in less than 150 trials in each session, in all three reward schedules (V2_HIGH, V2_MID and V2_LOW). This is referred as the ‘Control’ condition, green in Fig 4E-J.

Furthermore we simulated lesions of lateral and medial OFC in the model. Since the model generates a decision through at least one of the ‘limbic’ CBG loops and the other ASC and CBG_*pos*_ loop, even in case of a lesion to lOFC or mOFC, a valid decision should be made. We first describe the changes in the model with respect to each of the lesions and the corresponding results.

#### Medial OFC Lesion

A lesion of mOFC to the lmOFC model shown in Fig 3A makes the model slightly similar to the basic OFC model described in Fig 1B. The credit assignment still works by lOFC during the period after reward (Fig 1D, CBG, after ‘Reward’ phase), because the identity of the chosen cue maintained in lOFC is available for learning after the reward. However, in the absence of mOFC, the input to lOFC from the ongoing precise value comparison in mOFC is absent.

In all the control experiments, the model reached optimal performance within 150 trials. In the case of medial OFC lesions however, the performance was significantly impaired in the case of V2_HIGH scenarios, when V1 and V2 values were proximate. In the case of V2_MID and V2_LOW, the performance was observed to be similar to that of controls, except for a slight delay in reaching better performance. Such a normal performance in the case of V2_MID and V2_LOW can be attributed to the appropriate *credit assignment* by lateral OFC happening during learning. When the value difference is sufficiently large, and the credit correctly assigned to the correct choice, as the V2 anyway does not reward as much as V1, it is easily learned between the lOFC-CBG_*cue*_ synaptic connections and they can drive the decision without a precise comparison (Fig. 4 D-F).

#### Lateral OFC Lesion

One of the major changes in case of the lateral lesion is the credit assignment. In the control condition, when there is a reward delivered, the activation of the chosen cue in lOFC is active (Fig 1D, CBG, after ‘Reward’ until 2500ms). When there is no lOFC in the network, the association of current reward to only current choice can no longer be done. In this case, we still consider that the CSV for each cue is sent as an input to mOFC, because mOFC/vmPFC has been shown to receive projections from the ventral striatum [50, 51, 73], which is a crucial component of the CSV layer.

Striking of the observations, in the case when the difference between V1 and V2 was close, monkeys with mOFC lesions showed impaired performance while the controls and animals with lOFC lesions fairly performed well as V1 and V2 were distinct. Such an impairment argued for the role of mOFC to be more sensitive to the value difference between the options. Conversely, when the difference between the values was more, quite surprisingly animals with lOFC lesions were impaired whereas the controls and animals with mOFC lesions could steadily perform optimal choices. While it was an interesting observation to see how mOFC could not compensate even when the difference between the values is high (meaning it is an easy choice), it highlighted the role of lOFC in appropriate credit assignment, i.e. assigning the reward to the appropriate choice made in the current trial rather than to the previous or even the succeeding choice or even to the choice that rewarded the most historically.

In the case of lOFC lesions, the performance was affected in rather contrasting manner. Although eventually the performances reached near-optimal in all three cases of V2_HIGH, V2_MID and V2_LOW, the performance was sub-optimal for most of the earlier part of the sessions especially in the cases where the value difference was larger. This highlights the importance of lOFC in appropriately assigning the credit of reward to the correct option. Impairment of performance in the absence of lateral OFC in the case of V2_MID and V2_LOW was observed for the initial part of the session. This may be due to partial learning in the form of reward-based history maintained in medial OFC (as it was maintained when lateral OFC in intact) (Fig. 4 H-I).

## Discussion

We demonstrated the OFC on top of classical sub-cortical decision-making systems, with the descriptions of experimentally observed roles of its individual sub-regions. We explain the seemingly dissociated yet more complicated effects of the sub-regions of the OFC on the task performance depending on the task structure (value difference between the options). The OFC is clearly a crucial prefrontal region with heterogeneous representations and dynamics that result in complex behavior. Therefore clearly it is not a feasible idea to attempt a simplistic representation that relies on a unique way of information processing within the OFC, without implying several other brain regions that closely interact with the OFC during the behavior. Instead, we acknowledge the positioning of the OFC in the grand picture of several prefrontal and sub-cortical brain regions, as well as the heterogeneity within itself. Before attempting to model the possible mechanisms within the sub-regions of OFC in detail, it is crucial to build a framework that embeds a representation of environment in an embodied manner (with bodily needs and relevant behavior protocols for testing). Hence this work points towards the interest for modeling the dynamics of OFC as a part of a larger framework of related brain systems that the OFC interacts with, and the valuation systems it employs to guide decisions and learning. We chose one of the well-accounted frameworks of decision-making and reinforcement learning involving the BG (CBG loops) and complemented it with the specialized representations of the sub-regions of the OFC. We show that a simplistic model as in Fig 1B is sufficient for simple tasks. We further demonstrate that a more informed decomposed model (*lmOFC* model, Fig 3), while performing equally well on the simple tasks, also allows to study performance on more complex tasks in which the basic model cannot perform well.

### Choice

Although the primary role of lateral OFC has been implied to be appropriate credit assignment, here we use the fact that lateral OFC still plays an important role in driving the activities in the downstream BG loops with the dynamic subjective values added to the visual salience. Since the synaptic weights that are changed through learning are the ones connecting lOFC to the CBG_*cue*_ loop, the dynamics would passively favor a choice whose connection weights have been sufficiently learned. For instance, in the case of a lesion to mOFC (Fig 5), the initial decisions before any learning may be guided by lOFC through the CBG loop, randomly with the help of intrinsic noise. However, as in the case of Fig 5E,F where only cue rewards significantly more, the learning can rapidly increase the synaptic weights in the network corresponding to that cue and thus guide the subsequent decisions to that cue. This could be one reason why the performance slowly picked up towards the latter end of the trials, in the cases of V2_MID and V2_LOW in case of medial OFC lesions. Whereas in the case of mOFC lesion under V2_HIGH condition, since V1 and V2 almost similarly reward, even if appropriate credit assignment is done between the cues corresponding to V1 and V2, the network may not be able to definitively guide the decision necessarily towards V1.

### Learning

Learning in the system occurs at the level of both CSV (for expected values) and cortico-striatal synapses. The learning that occurs at the level of cortico-striatal synapses indirectly represents the reward contingencies of stimuli in terms of their probability. One of the possible motivations behind multiple learning mechanisms in the system is the feasibility of a shift of control from the value-comparison based processes in the mOFC/vmPFC at the beginning of the trials to a faster, network strength based decision through the lOFC-BG loops driven by the learned connection weights, in the trials after substantial learning. Such a distinction was reported where activities in vmPFC were more remarkably distinct between more deliberative situations with slower reaction times as opposed to trials towards the end of the experiment or even no-brainer trials (highly probable high reward versus the opposite). Moreover, the involvement of value-difference signal in vmPFC consistently decreased towards the later trials of the task [74]. However, it is important to note that, in a different formal description, it has been highlighted that ventrolateral PFC (vlPFC) encodes the *Availability* (probability) of rewards whereas the OFC was shown to encode the *Desirability* (palatability) of rewards [75]. However it was shown activity in medial and lateral orbitofrontal cortex, extending into vmPFC, was correlated with the probability assigned to the action actually chosen on a given trial [70].

#### Lateral and Medial in Learning and Choice : Dissociation or Interaction ?

What is important to note here is that the dissociate effect observed does not necessarily imply that both the sub-regions have dissociate roles in decision-making and learning, as recently suggested [56]. The dissociation might be observed in terms of the anatomical connectivity in the sense that lateral OFC predominantly receives inputs from sensory regions about external environment whereas medial OFC has more inputs from internal bodily states and visceral responses. But we argue that seeming dissociation in terms of internal processes within these sub-regions is a possible network effect as a result of their temporal dynamics. Because, as we have shown here, both the sub-regions are involved in the circuitry that is capable of both guiding decisions and learning from outcomes. Albeit, the dissociation might be apparent because of the fact that each of them might have access to different information about the state of the environment and exert control at a different stage of the behavior.

### Extensions

The task structures used in this work are related to only one type of outcome and assuming the motivational value of the outcome is always non-zero. And ventral striatum particularly plays an important role in learning action-values as well [4]. By incorporating computational accounts of emotional value learning in BLA, and taking the internal motivation into account through ventral striatum and lateral hypothalamus, experimental results of the role of OFC in paradigms like Reinforcer Devaluation [28, 42, 52] can be explained. Possibly, interesting findings like the one in Reversal Learning paradigm can be explored where neither lateral nor medial lesions of OFC do not affect the behavior whereas the lesion of OFC as a whole affects. As mentioned earlier, the OFC has been proposed to represent a cognitive map of task space [9] and to encode the *value* of the offered and chosen goods [10, 11]

#### State/Task space representation in the OFC

The OFC has been proposed to encode the task states and represent a cognitive map of this task space [9]. It has also been shown that OFC lesions in animals cause deficits in acquiring information about the task [27, 76]. In this work, related to the simple 2-arm bandit task done under a DIFFICULT condition (Fig 2D, red), the model fails to perform as good as in the EASY condition. However, with a slight change in the task structure, the model can be shown to perform better. That is, instead of presenting the same pair of shapes, imagine there are four possible shapes and the 6 possible pairs are presented for choice pseudo-randomly. Even if the reward probabilities of both the shapes used in Fig. 2A remain the same, besides two other possible options, the options with which these cues are presented change. This can be due to the modified learning rates because of the state-change across each trial (because no pair is presented consecutively). Alternatively it can also be remarked that, even though the value of the first two shapes did not change, the overall value of each trial or that of the entire task has changed in the presence of other options. These two factors can be hypothesized to be represented in terms of state prediction errors and value difference signals in lateral and medial OFC respectively.

#### Temporal Dynamics : Delayed presentations, Opportunistic behaviors

One of the clear limitations of the model presented in this work is that the temporal dynamics at various stages of decision-making processes with respect to the discussed sub-regions of OFC is not entirely accounted for. Particularly an intracranial EEG recordings of OFC in humans showed that lOFC was encoding experienced value in the reward delivery phase [77]. This model needs to extended further by incorporating the cortico-cortical interactions, within the subregions of OFC as well as other related prefrontal regions like the ACC which are believed to be rather interactive in their roles in behavior [78]. Furthermore, the role of any possible interaction between both the sub-regions of OFC, given their connectivity through the medial orbital sulci [46], has not been explored much. However, even if it is the case that the lateral OFC represents identity specific rewards and vmPFC represents general, scaled reward signals, it is unclear how these two signals could be linked to sub-serve goal-directed behavior. To this extent, there is not much evidence except one study that showed the functional connectivity was predictive of satiety related changes in choice behaviour [79]. Similarly, the role of VS also becomes crucial in serving such value signals that combine both value and internal motivation before the comparison processes in the mOFC/vmPFC.

## Conclusion

The anatomical description implicitly highlights the difference in the accessibility of lateral and medial OFC for experimentation procedures. The same applies to imaging studies, which happen to be a major contribution of studies in humans, that the regions highlighted by BOLD signals cannot be precise enough within the scope of subregions. Another challenge is the homologies of the OFC among humans, nonhuman primates and rodents. Since a good part of the literature on OFC is almost equally contributed by the studies in all three species, it would be major task at hand to be wary of the similarities and the differences among what is defined as OFC in each of these species. Functionally, it is also not straightforward to identify whether a difference in an ability of one species (say humans) to demonstrate a faculty and that of another (say rats) is a difference in kind or a difference in degree. Although, it would be fairly possible to extend the conclusions from one species to another depending on what is being studied (for example, the findings related to action-outcome contingencies or basic behavior in rats might extend well to primates beyond which more flexible representations might emerge).

### Challenges in studying the representation and mechanisms in OFC

As far as the interest in the dissociate contribution of subregions of OFC is concerned, there are not so many experimental evidences that could establish a double dissociation between different subregions [28, 42, 43, 80]. Importantly, it has been found out that both lateral and medial regions of OFC represent the perceived value of task events, albeit with different levels of participation in different task settings [42]. It is generally single-neuron recording studies or lesion studies in macaques or rats [81] predominantly on the better accessible lateral OFC than the medial OFC, and BOLD signal correlation from fMRI studies in humans. Few behavioral studies on frontal damage patients have also discussed separate roles of OFC and vmPFC [4, 26, 82, 83]. However, owing to the different techniques and methodologies used, there are few inconsistencies where both lateral and medial regions were exclusively implied for signals during the anticipation of rewards [84]. Another interesting theory about the dissociating role of lOFC and mOFC that probably requires closer look is that while mOFC might represent the values of options in a context where there is no choice to be made, lOFC doesn’t represent the value in a choice-free context [80].

Notwithstanding some complementary [85, 86] as well as contrasting [87, 88] findings, separable representations of absolute values and relative values seem to be a key dissociation between the medial [5, 19, 89, 90] and lateral [91] OFC. Future studies that could effectively dissociate the factors such as salience of the stimuli, internal motivation and thereby investigate the difference between absolute and relative value encoding, should provide more insights into whether lateral and medial OFC play dissociate role to represent them.

Well within one species, reported counter-intuitive findings also might encourage the need to dissociate the roles of lateral and medial OFC. Quite often the ambiguity arises from the task structure which doesn’t sufficiently dissociate closely related aspects of a certain behavior. In the few studies that have described the nature of dissociation between lateral and medial OFC, certain results were difficult to explain and there appears to be several possibilities for such results. For instance, when it is observed that neither of the individual lesions of lateral and medial OFC impair the animal in Reversal Learning while the lesion of OFC as a whole does [83, 92], it could be possibly mediated by other sub regions of OFC (central, anterior or posterior) or there might very well be a possible mechanism through which they partially compensate for one another interacting with other parts of the brain. In consecutive reinforcer-devaluation tests, while monkeys with lOFC lesion showed significant impairment, the ones with mOFC lesion were less consistent in their choices across sessions contrasting overall performance similar to that of lateral lesions immediately after the lesion, and with that of the controls in the later sessions after the lesion [28].

## Materials and methods

We use a neuro-computational connectionist modeling approach to highlight the organization of subsystems that drive decision-making and learning. The subsystems are built with simplified representations of the experimental findings related to the roles of lateral and medial OFC. The comprehensive model accommodates representations of a fairly complete set of phases involved in decision-making [8]. Besides the component of OFC, the information processing in the model for decision-making and learning also involves minimized yet biologically plausible representation of several sub-cortical mechanisms involved. However, we emphasize more on the contributions of the lateral and medial OFC, maintaining the generic nature of rest of the mechanisms. We first introduce the kind of tasks on which the model is tested, that would demonstrate both the dynamics of a decision as well as the progression of behavior through the task. Then the major computational aspects of the model will be presented, referring to the representations required for the tasks described.

### Thalamo-Cortical Basal Ganglia (CBG) Loops

The fundamental decision making networks and learning in the system are implemented in the form of schematic thalamo-cortical basal ganglia (CBG) loops which process the information of the cues, estimated values of the cues and the actions required to select the cues. The core action selection mechanism between multiple options - cues or actions, is implemented using an architecture of the basal ganglia (BG) similar to that has been described in classical descriptions of pathways in BG (summarized well in [93]). Thus we define a CBG loop as a thalamo-cortical BG loop, an example of which is described in S1 Fig. The architecture presents a general idea of the connectivity between the input structures of BG - Subthalamic Nucleus (STN) and Striatum (STR) and the output structures - Globus Pallidus pars Interna (GPi) and Substantia Nigra pars Reticulata (SNr).

We implement computational model of parallel loops of three kinds, which were originally described as : *limbic, sensori-motor* and *associative* [94]. The *limbic* loops originate in the orbitomedial prefrontal cortex (generally comprised of OFC and ACC, through amygdala, hypothalamus and the subdivisions of VS (nucleus accumbens) and end back in the medial PFC. The *limbic* loops, besides processing external information, are based on interoceptive information. They are organized around the selection of the goal of the behavior, according to its motivational value, in response to perceived needs or according to its hedonic value. Individual sub-regions of medial prefrontal cortex form the feedback loop through different nuclei of VS [95, 96]. It is these *limbic* loops that we focus on in this work, specifically emphasizing the role of the OFC and possible dissociate roles of its sub-regions, since lateral and medial OFC also are part of this network of parallel loops. For example, in a 2-arm bandit task shown in Fig 1A, the cues (shapes) that are presented in each trial are represented within these *limbic* loops (CBG_*cue*_ in Fig 1A). The information about the position of the cue (thus the required action to select the cue) is represented in the sensori-motor loops (CBG_*pos*_ in Fig 1A), from the regions in Parietal Cortex to form the feedback loop through the dorsolateral striatum (DLS) [97, 98]. The lateral prefrontal cortex (lPFC) forms an associative loop with the dorsomedial striatum (DMS), receiving multimodal information from the associative regions of the posterior cortex (ASC in Fig 1A). The combined information of which cue is present in which position, which solves the binding-problem, is represented in the lPFC and DMS [62].

The population dynamics and the learning mechanisms described below have been adapted from similar works before on the thalamo-cortical BG loops [59, 60].

### Population Dynamics

All the structures within a CBG loop are implemented as populations that are part of a recurrent neuronal network, with neuron units of similar dynamics. The dynamics of a neural population unit is described in equation 1, similar to previous computational accounts of decision-making in the BG [63]. Assuming each population unit represents an ensemble tuned towards a particular option : *I_ext_* is the external input representing the salience of the option, *I_s_* is the input to the unit from its connections (synaptic input) and *τ* is the decay time constant of the synaptic input and *V* is the resultant activity of the unit. External input, *I_Ext_* is provided only in cortical structures (for the other structures *I_Ext_* = 0), and T is the threshold of a neuron, depending on the population. Also, symmetry breaking is generated by Gaussian noise *δ* to the activity of each ensemble at each time step.

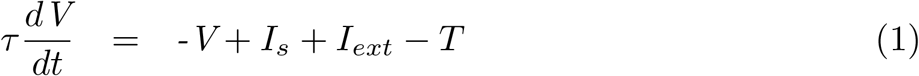

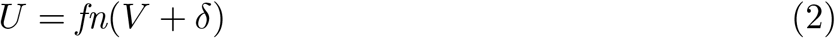

The activation function *fn* in Eq. 2 is the same for all the structures within a CBG loop and it is a clamping function, except for the striatal structures. The activation of striatal populations, due to their neuronal properties [99–101], can be obtained by applying a sigmoidal transfer function to the activation of CTX-STR inputs in the form of the Boltzmann equation S1 Equation.

The synaptic input to a unit *j*, 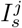, which is the input as a result of the connections from units of other structures (say *i*), depends on the connection weights (*w_ij_*) between units *i* and *j*, as shown in the equation 3. Except the synaptic connections that can be learned, the rest remains to be constant connection weights chosen at the beginning, within the range of 0.25 and 0.75, generally chosen around 0.5.

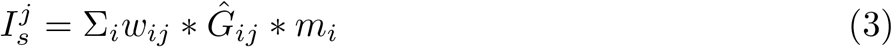

Also, there is a fixed gain parameter that characterizes the strength of interaction between the two populations to which *i* and *j* belong. For example, for any pair of connections *ij* between CTX(*i*) and STR(*j*), the gain 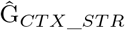 is fixed. A positive or negative 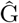 defines the connection as excitatory or inhibitory respectively. In the “direct” pathway, as a result of two inhibitory and one excitatory connection, it is referred as a positive feedback loop. In the “hyperdirect” pathway, as a result of two excitatory and one inhibitory connection, it is referred as a negative feedback loop [63].

### Learning

The connections between the OFC and the CBG_*cue*_ loop in the basic model (Fig. 1B) are modifiable. Similarly, after the model is changed to lmOFC model (Fig. 3), the connections between lOFC and its CBG loop, as well as mOFC and ASC are modifiable. The modifiable connections between lOFC and its CBG loop are One-To-One between cue populations in lOFC and cue specific populations in the structures of CBG loop. Whereas the modifiable connections between mOFC and the ASC network are One-To-All i.e, from each cue population in mOFC to all four position populations possible. After every decision and verifying the outcome, the weights are updated. Like in the previous models, all synaptic weights are initialized to 0.5 (SD, 0.005). The weight update term Δ*W_t_*, is calculated as a function of reward prediction error (RPE), which is believed to be signalled by dopamine at the level of cortico-striatal synapses. However, it was specifically found that striatal neurons involved in cortico-striatal synapses show long term potentiation (LTP) and long term depression (LTD) with respect to positive or negative prediction error, respectively ([102]). RPE precisely is the difference between the perceived reward value and the expected reward value. In the model, similar to a standard critic-learning RL framework, expected reward values of each stimulus population are maintained and updated.

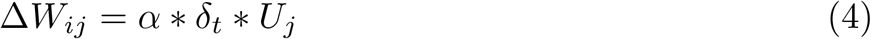

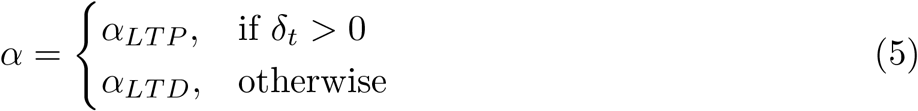

The RPE, *δ_t_* is calculated using a simple critic learning algorithm given below.

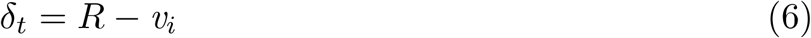

where R, the reward, is 0 or 1, depending on whether a reward was given or not on that trial. After the Δ*W_t_*, is calculated, the synaptic weights are updated according to S2 Equation. And upon weight changes, to make sure the weights stay within the initial bounds, every weight update is followed by a normalization of weights (S3 Equation). *v_i_* is the CSV of the cue represented by neuron *i* in the CBG. The CSV of the chosen cue is then updated by :

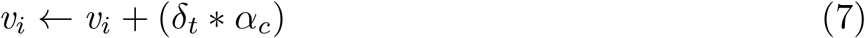

where *α_c_* is the critic learning rate and is set to 0.025 and *α_LTP_* and *α_LTD_* are set to 0.004 and 0.002 respectively.

### Current Subjective Value (CSV)

OFC is known to represent a current subjective value (CSV) of a stimulus with respect to the body’s internal state (like satiety or desirability of the outcome the stimulus announces). Two primary brain structures that crucially involve with the OFC in this regard are : the amygdala and the ventral striatum (VS). The basolateral amygdala (BLA) has been shown to interact with the OFC and update its stimulus-outcome associations and hence the subjective value of a stimulus [53, 54]. On the other hand, the ventral striatum was found to represent a unified quantity as a combination of subjective value and internal motivation using different kind of neurons [55]. Several computational accounts have explained possible implementations of such representations [103–107]. A much detailed representation and role of ventral striatum and its distinct relation to lateral and medial OFC also could be a key factor to study [108–111].

## Supporting information

**S1 Fig High level architecture of Thalamo-Cortical-BG Loops in primates.** Classic BG connectivity : STN and STR as inputs, GPi/SNr as outputs. GPi: Globus Pallidus pars Interna; SNr: Substantia Nigra pars Reticulata; STN: Subthalamic nucleus; STR: Striatum. The direct pathway from the prefrontal cortex (PFC, here used generally, including OFC) via STR to GPi, and the hyperdirect pathway from CTX via STN to GPi. It has to be noted that this is only one of several possible interpretations of action selection mechanism within BG, as it sufficiently explains using one excitatory and inhibitory pathway. Several other interpretations exist, for instance, an indirect pathway, which are not considered in this work. Indirect pathway, involving STN, GPe (Globus Pallidus pars externa) and STR, is also a part of “classical” view of CBG network [112–114]. Image re-illustrated, inspired from [93]

**S1 Appendix.**

**S1 Equation. Sigmoidal transfer function**

**S2 Equation. Oja weight update rule**

**S3 Equation. Weight normalization**

**S1 Table. Parameters of CBG loop structures**

## Footnotes

1 What was referred to as OFC in this work actually recorded from the lateral areas of the OFC

